# The fetal specific gene *LIN28B* is essential for human fetal B-lymphopoiesis and initiation of KMT2A::AFF1 infant leukemia

**DOI:** 10.1101/2024.09.18.613730

**Authors:** Rebecca Ling, Thomas Jackson, Natalina Elliott, Joe Cross, Lucy Hamer, Arundhati Wuppalapati, Alastair Smith, Catherine Chahrour, Okan Sevim, Deena Iskander, Guanlin Wang, Siobhan Rice, Sorcha O’Byrne, Joe Harman, Bethan Psaila, Rhys Morgan, Irene Roberts, Thomas A. Milne, Anindita Roy

**Author notes:** **CORRESPONDING AUTHOR:** Anindita Roy, Department of Paediatrics, University of Oxford, John Radcliffe Hospital, Oxford, UK, OX3 9DS.

## Abstract

Infant ALL (iALL) is initiated *in utero*, most often by rearrangement of the *KMT2A* gene (*KMT2Ar*). It carries a very poor prognosis despite a lack of additional oncogenic driver mutations common in childhood ALL. Here, we aimed to identify specific properties of human fetal hematopoietic stem/progenitor cells (HSPC) that promote leukemic transformation in *KMT2Ar* iALL using molecular, functional and *in vivo* assays. First, by comparing transcriptomes of human fetal HSPC to adult HSPC we derived a fetal-specific gene signature and identified the fetal oncogene *LIN28B* and its downstream effectors among the top hits. These genes were also expressed in iALL. Functional assays revealed that *LIN28B* was essential in human fetal liver (FL) CD34+ cells to maintain proliferation and stemness, and support B- and NK-lymphopoiesis. To interrogate the role of *LIN28B* in iALL, we utilised a human FL-derived CRISPR-Cas9 KMT2A::AFF1 model. In this model, *LIN28B*-expressing leukemias were more proliferative *in vitro* and *in vivo*, with this advantage being lost upon *LIN28B* knockdown. Mechanistic studies showed that LIN28B acts by stabilizing key early B-lymphoid genes, epigenetic regulators, and cell cycle and anti-apoptotic genes. Finally, In the absence of *LIN28B*, human FL CD34+ cells fail to transform upon induction of KMT2A::AFF1 translocation. Thus, *LIN28B* has an essential role in normal human fetal B-lymphopoiesis, and is necessary for the initiation of *KMT2A::AFF1* iALL in fetal cells in the absence of co-operating mutations. It has a role in making leukemias more aggressive, suggesting it is a potential target in *LIN28B*-expressing leukemias.

## INTRODUCTION

Infant Acute Lymphoblastic Leukemia (iALL) is a biologically distinct disease from childhood ALL (chALL)(1). Clinically, it is characterised by hyperleukocytosis, hepatosplenomegaly and central nervous system involvement at diagnosis, and a very poor prognosis despite several decades of international collaborative efforts to improve outcomes(2-4). Data from recent treatment protocols using targeted immunotherapies are promising, but still preliminary(5-8). The leukemia is predominantly B-lymphoid, with an immature CD19+CD10-immunophenotype, often with aberrant expression of myeloid markers(9, 10). Frequently, patients have an initial period of response to ALL therapy followed by a very early relapse, with >90% of these occurring within one year of starting treatment(2, 4). In a minority of cases, there is a lineage switch at relapse to myeloid leukemia, the likelihood of which may be higher after CD19-directed immunotherapies(11-14). The molecular landscape of iALL is dominated by *KMT2A* gene rearrangements which are seen in 75% of the cases, the commonest of which results in the *KMT2A::AFF1* gene fusion(2, 15). The genetic landscape of iALL is otherwise silent, with secondary mutations being rare, and when present they are subclonal/non-essential(16-18). Importantly, *KMT2Ar* chALL is less aggressive and more responsive to therapy than *KMT2A::AFF1* iALL(19-22). It is difficult to understand why a seemingly genetically simple, ‘one-hit’ B-ALL is so much more aggressive than the cytogenetically matched *KMT2Ar* chALL. While there has been much debate about a different and more immature cell of origin in iALL driving these features, the developmental stage specific origins of the disease, and how the cell of origin might contribute to leukemogenesis remains poorly understood.

It has been known for a while that iALL is initiated *in utero* when *KMT2Ar* occurs in a fetal progenitor cell(23-28). This single event seems sufficient to cause the rapid onset of an aggressive ALL within the first few months of life. Whilst a number of chALL gene fusions (ETV6-RUNX1, BCR-ABL, TCF3-PBX1, TCF3-ZNF384 and KMT2Ar)(26, 28-32) also have *in utero* origins, the onset of these leukemias is much less rapid and require a ‘second hit’ of additional co-operating mutations. This suggests that specific properties of human fetal hematopoietic stem and progenitor cells (HSPC) may provide the appropriate cellular context that promotes leukemic transformation in *KMT2Ar* iALL(33). Supporting this hypothesis are data from several studies that have mapped the transcriptomic, metabolic and proteomic profile of *KMT2Ar* iALL to fetal HSPC(34-39). However, the strongest evidence for the necessity of a human fetal cell context for initiation of *KMT2Ar* iALL comes from the inability to generate an accurate animal model of KMT2A::AFF1 iALL till the fusion gene was expressed in human neonatal and fetal HSPC(40-42). This led us to hypothesize that the expression of fetal-specific genes provides the permissive context for the initiation of KMT2A::AFF1 iALL and that their ongoing expression impacts the disease phenotype.

To investigate this, we first compared the transcriptomes of human fetal HSPC to adult HSPC to derive a fetal-specific gene signature and identified the fetal-specific oncogene *LIN28B* and its downstream effectors *IGF2BP1, IGF2BP3* and *HMGA2* (43, 44) amongst the top hits (Suppl Fig 1a). *LIN28B* is aberrantly expressed in many cancers(44-48), and several of its target genes, such as *IGF2BP1* and *IGF2BP3*, are known to stabilise key KMT2A::AFF1 target oncogenes, such as *HOX* genes(49, 50). *HMGA2* is a chromatin remodelling factor which enhances cell proliferation in KMT2A::AFF1 ALL(51). In addition, in mice, *Lin28b* and its partner gene *Igf2bp3* have been directly implicated in the fetal HSC phenotype of proliferation, self-renewal and pluripotency, and specifies fetal-like lymphopoiesis(43, 52, 53).

**Figure 1.**
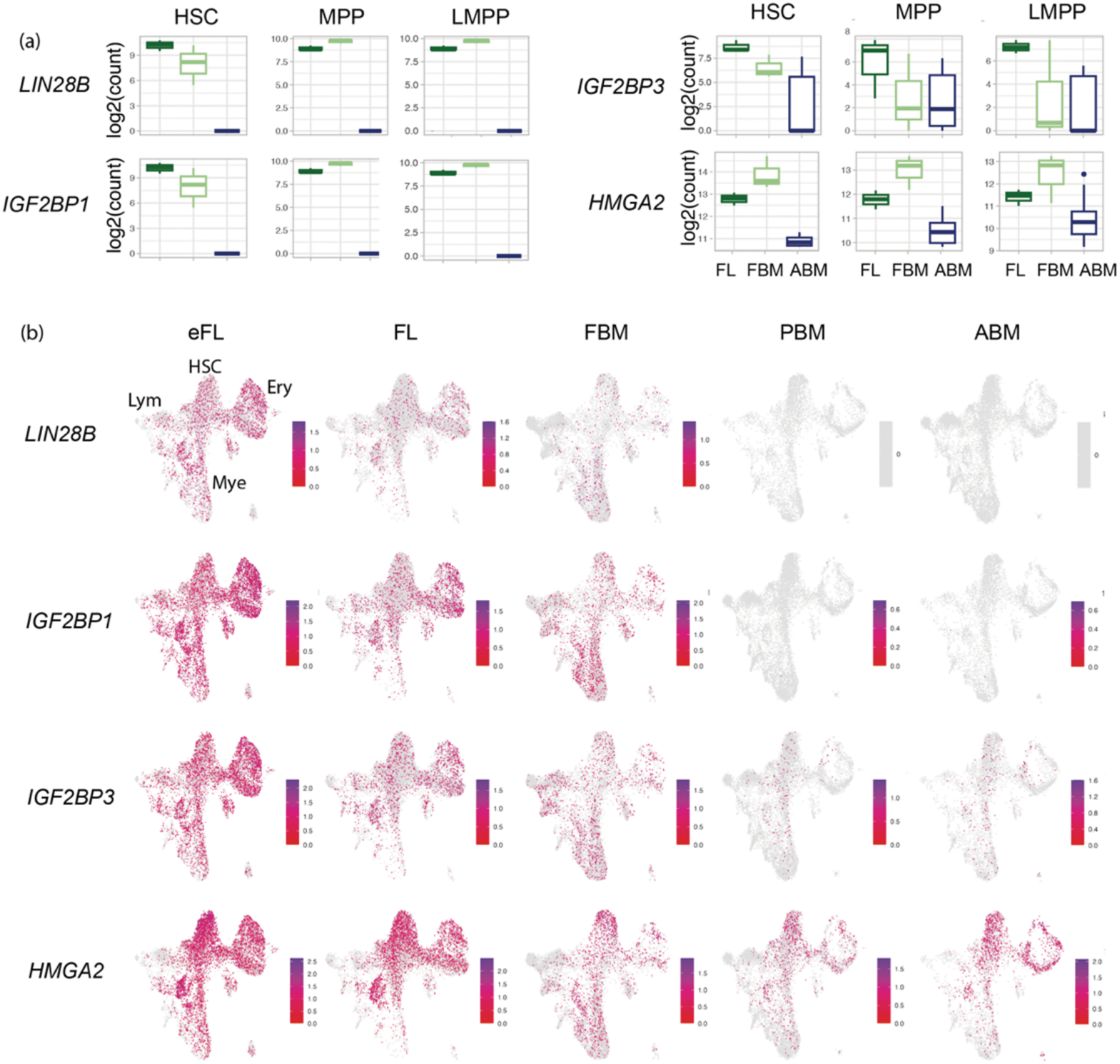
(a) Bulk RNAseq data from sorted HSC, MPP and LMPP from FL (n=3), FBM (n=3) and ABM (n=3) showing expression of *LIN28B* and key partner genes. Data presented as log2 normalised counts (b) scRNAseq UMAP for Lin-CD34+ HSPC from (55), showing expression of *LIN28B, IGF2BP1, IGF2BP3* and *HMGA2* across ontogeny in first trimester FL (early FL: eFL), paired second trimester FL and FBM, paediatric BM (PBM) and ABM, total = 57489 cells. (Lin: lineage markers: CD2,3,14,19,56,235a; HSC; hematopoietic stem cell cluster; Ery: erythroid progenitor cluster; Lym: early lymphoid progenitor cluster; Mye: myeloid progenitor cluster)

Here, we show that *LIN28B* and its downstream effectors are not only highly enriched in human fetal HSPC, but also expressed in iALL. To test the hypothesis that *LIN28B* is essential for iALL initiation by KMT2Ar and the potential mechanisms responsible, we used a human FL-derived CRISPR-Cas9 KMT2A::AFF1 model and show that in human FL CD34+ cells, *LIN28B* is essential for normal B- and NK-lymphopoiesis and, in turn, for leukemic transformation of fetal HSPC by KMT2A::AFF1. These data provide the first evidence of a role for LIN28B in human fetal B-lymphopoiesis and infant leukemia initiation, and highlights possible mechanisms that could be targeted in this aggressive disease.

## RESULTS

### *LIN28B* expression is highest in human fetal HSC, MPP and LMPP

*LIN28B* is known to be highly expressed in fetal tissues, including primitive hematopoietic progenitors(54) but a detailed analysis of its expression in human fetal HSPC subpopulations has not been performed. To understand the physiological role of *LIN28B* in human fetal hematopoiesis, we first characterized its pattern of expression in normal fetal HSPC using mini-bulk RNA-sequencing data from human fetal bone marrow (FBM)(35) and FL(36), and single-cell (sc) qPCR data from FBM and adult BM (ABM)(35). In fetal HSPC, *LIN28B* expression was highest in HSC (Lin-CD34+CD38-CD45RA-CD90+), MPP (Lin-CD34+CD38-CD45RA-CD90-) and LMPP (Lin-CD34+CD38-CD45RA+) compared to more differentiated B-lymphoid progenitors (Suppl Fig 1b). Expression in HSPC from ABM was undetectable by sc qPCR (35) (Suppl Fig 1c). Differential gene expression analysis of bulk RNAseq data from fetal (35, 36) and ABM HSC, MPP and LMPP (Suppl Fig 1d) showed that in addition to *LIN28B*, major *LIN28B* pathway genes, including *IGF2BP1* and *HMGA2* were upregulated in FL and FBM HSC, MPP and LMPP compared to ABM counterparts (Fig 1a, Supplementary Table 1). Analysis of our previously published sc transcriptomic data (55) revealed that expression of *LIN28B, IGF2BP1, IGF2BP3* and *HMGA2* was highest in first trimester ‘early’ FL (eFL) Lin-CD34+ cells and decreased through ontogeny. *LIN28B* and *IGF2BP1* are fetal specific genes, with their expression being completely absent in paediatric BM (PBM) and ABM HSPC. Within the Lin-CD34+ fraction, *LIN28B* pathway genes have heterogeneous expression, and are most highly expressed in HSC/MPP and erythro-myeloid progenitor clusters, with almost no expression in early lymphoid progenitor cells (Fig 1b).

### *LIN28B* is necessary for human fetal B-lymphopoiesis

To investigate the functional role of *LIN28B* in normal fetal hematopoiesis, we used lentiviral transduction of *LIN28B* shRNA into FL CD34+ cells (4 biological samples: 12, 17, 20 and 21 pcw) (Suppl Fig 2a,b) and measured the impact on gene expression and functional output *in vitro* and *in vivo*. Knock down (KD) of LIN28B achieved >3-fold reduction in *LIN28B* expression compared to scrambled RNA (scr control) (Suppl Fig2c,d). RNAseq analysis showed 1306 DEG (log2FC>0.5, padj<0.05) with downregulation of a fetal lymphoid geneset (NES -1.94, padj 0.0017)(56). Of note, B-, NK and T-lineage gene sets were all downregulated, including : *IKZF1, CD79A, IRF4, FLT3, POU2F2*; *IL2RB* and *CD69*; and *GATA3* and *ETS1* respectively (Suppl Fig 2e)(56). There was a concomitant upregulation of a fetal myeloid geneset (NES 2.16, padj 0.0027)(56), including *CEBPB, FGL2, CD46, CD55, CTSB* and *ITGA2B* in *LIN28B* KD FL CD34+ cells (Suppl Fig 2f)(56). There was a downregulation of HSC genes (57) including *CD44, KDM2A, HLF, MLLT3, FLT3* and *MEIS1* (Suppl Fig 2g). Genesets related to cell proliferation, MYC targets (padj (3.7×10^−6^) (*MEIS2, MCM6, ATF6, RPS6KAS, G3BP1)* and *MYB* target genes (padj 9×10^*−*5^) were also downregulated. Together these data suggest that *LIN28B* KD causes a molecular reprogramming of FL CD34+ cells towards myelopoiesis at the expense of lymphopoiesis, with concomitant downregulation of stem cell and proliferative signatures.

**Figure 2.**
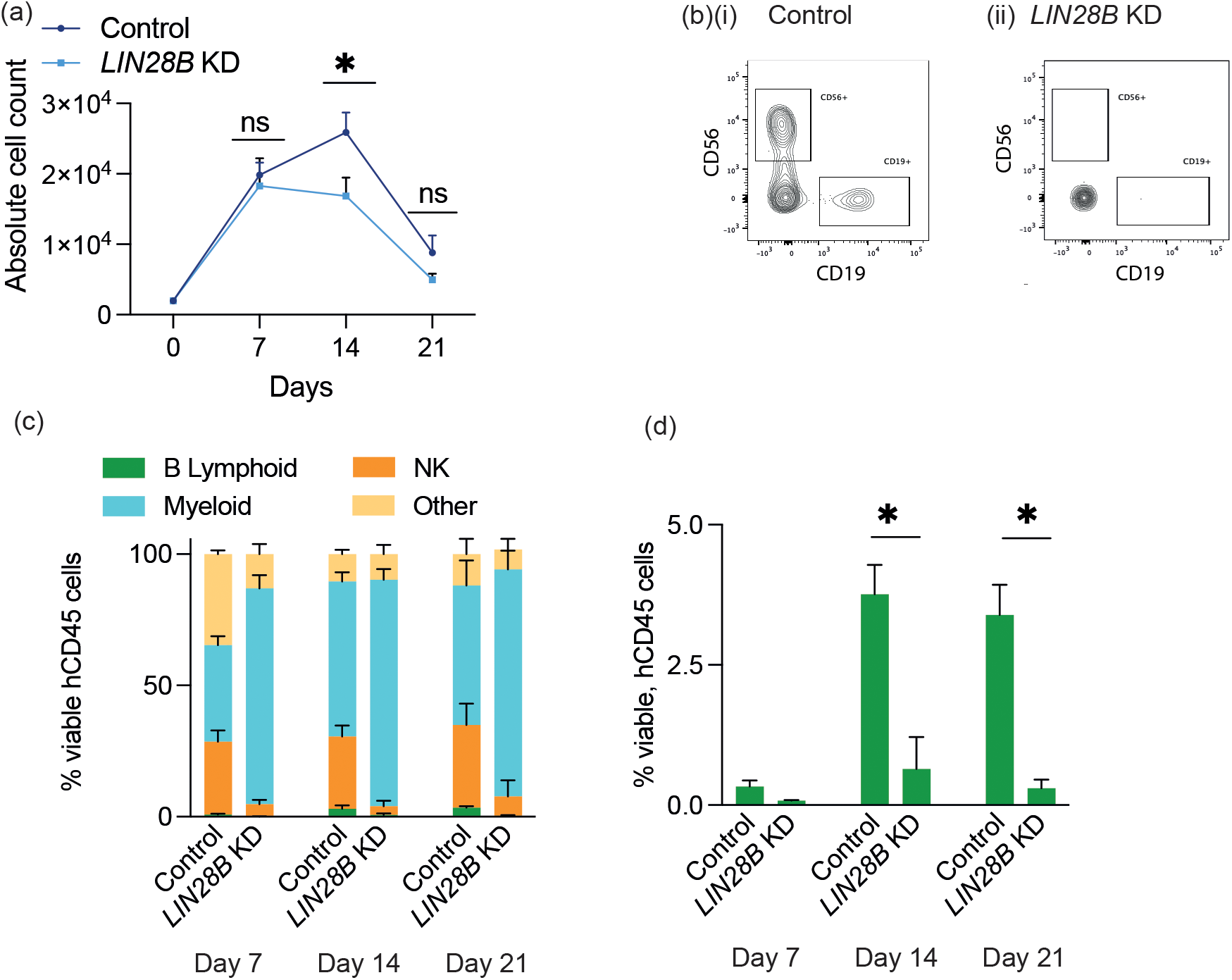
(a) Proliferation of control or *LIN28B* KD FL CD34+ on MS5 stroma over 21 days (b) Representative flow cytometry plots of cells harvested at day 14 from (i) control and (ii) *LIN28B* KD wells, showing markedly reduced CD19 and CD56 output from *LIN28B* KD FL CD34+ cells. (c) Graphs showing proportion of B-lymphoid (CD19+), myeloid (CD11b/CD14/CD16+) and NK cells (CD56+) at weekly intervals for control or *LIN28B* KD FL CD34+ cells (data shown as % of viable hCD45 cells, mean+SEM) (d) B-lymphoid output at Day 7, 14 and 21 from control or *LIN28B* KD FL CD34+ on MS5 stroma (data shown as % of viable hCD45 cells, mean+SEM. *p<0.05).

To test the functional relevance of these findings, we assessed the growth and lineage output of *LIN28B* KD and control FL CD34+ cells using an MS-5 stromal co-culture system. Total cellular output was reduced by day 14 of culture from *LIN28B* KD FL CD34+ cells (Figure 2a), particularly of B-lymphoid (p = 0.024) and NK cells (p = 0.0002) in tandem with an increase in myeloid output (p= 0.0025) (Figure 2b-d). These data support the functional importance of the gene expression changes after LIN28B KD and suggest that, as in mice(52, 53), *LIN28B* plays an important role in human fetal B- and NK-lymphopoiesis.

### *LIN28B* is required for *KMT2A::AFF1*-induced leukemic transformation of human fetal HSPC

Studying the initiation of iALL requires a relevant model system where one can induce the leukemia. We have previously shown that CRISPR-Cas9 editing to induce a *KMT2A::AFF1* translocation in normal human FL HSPC (^CRISPR-*KMT2A::AFF1*^ALL) accurately recapitulates iALL(40). The ^CRISPR-*KMT2A::AFF1*^ALL model was utilized to test whether *LIN28B* is necessary for leukemia initiation. LV KD of *LIN28B* and CRISPR-Cas9 editing to induce *KMT2A::AFF1* were performed sequentially in FL CD34+ cells (n=3 biological samples, pcw = 15, 17, 21) (Figure 3a). The presence of the *KMT2A::AFF1* fusion gene was confirmed in both control and *LIN28B* KD cells by digital droplet PCR (ddPCR) with an editing efficiency similar to un-transduced FL CD34+ cells (1-2%) (Figure 3b). Leukemic transformation *in vitro* was assessed by seeding on MS5 stroma and defined by serial measurement of cell numbers, immunophenotyping and morphology. *In vitro* evidence of transformation occurred in 5/19 (26.3%) ^CRISPR*-KMT2A::AFF1*^control FL CD34+ wells compared to 0/18 wells seeded with ^CRISPR*-KMT2A::AFF1*^*LIN28B* KD FL CD34+ cells (Figure 3c). There was a marked difference in cell proliferation between transformed (n=5) and untransformed (n=14) ^CRISPR*-KMT2A::AFF1*^control wells (p=0.000056). ^CRISPR-*KMT2A::AFF1*^*LIN28B* KD cells proliferated at a similar rate as untransformed ^CRISPR-*KMT2A::AFF1*^control cells (p= 0.68) (Figure 3d). By day 21, >90% of the cells in transformed ^CRISPR-*KMT2A::AFF1*^control wells were CD19+ whereas, non-transformed ^CRISPR*-KMT2A::AFF1*^control wells showed the expected multilineage output of B-lymphoid, myeloid and NK cells, and ^CRISPR*-KMT2A::AFF1*^*LIN28B* KD wells were predominantly myeloid, with <5% CD19+ cells (Figure 3e) consistent with the data in Figure 2.

**Figure 3.**
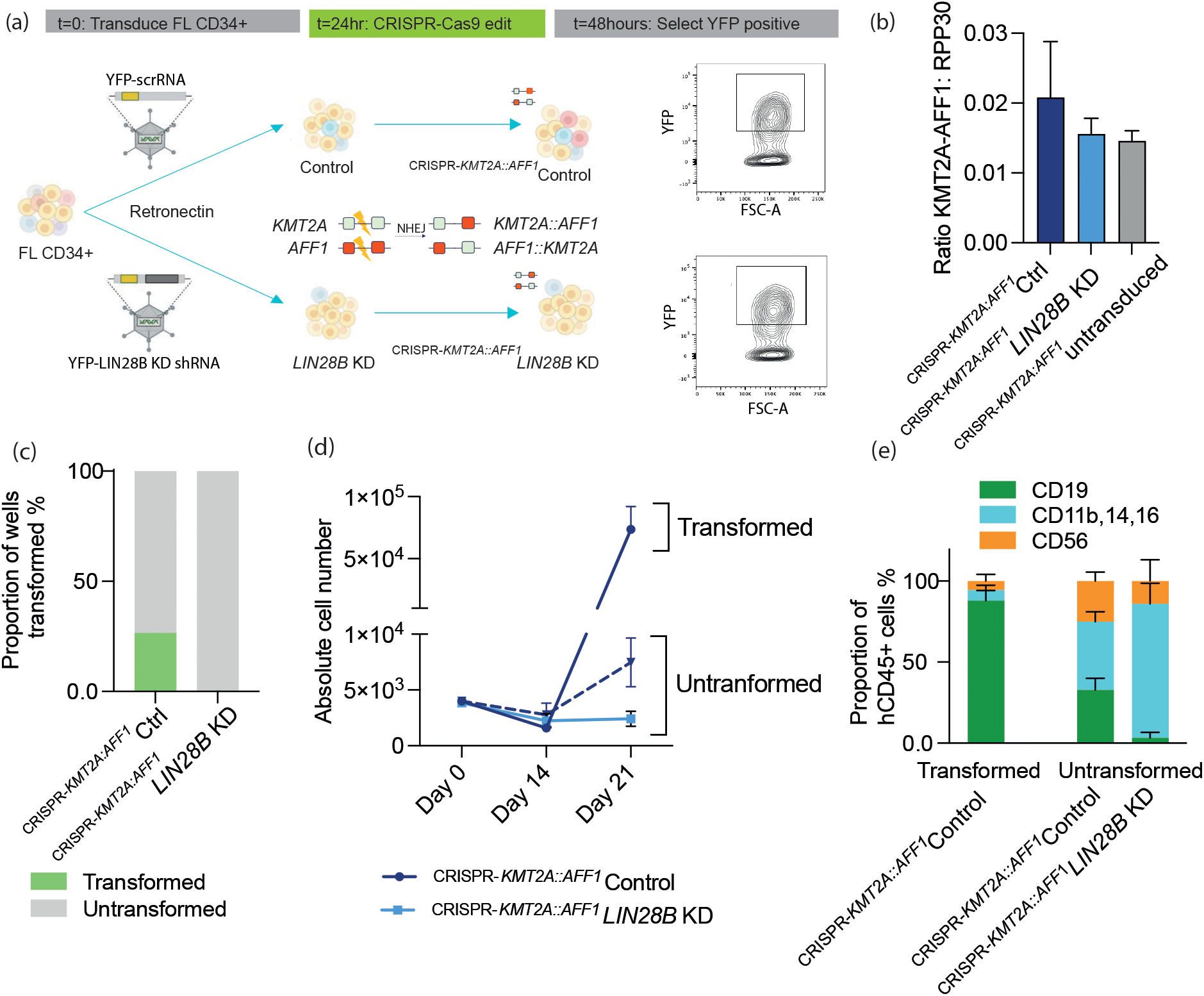
(a) Schema for LIN28B KD followed by CRISPR induced *KMT2A-AFF1* initiation of leukemia in human FL CD34+ cells. (b) *KMT2A::AFF1* fusion gene quantification by ddPCR in control and LIN28B KD FL CD34+ cells (n=3) and untransduced FL CD34+ cells (n=3). (c) Leukemic transformation *in vitro* expressed as proportion of wells that transformed for control (Cntrl) FL cells (n= 20 wells, 4 biological replicates) and LIN28B KD FL cells (n=18 wells, n= 4 matched biological replicates) (d) Cell proliferation depicted as absolute numbers of human CD45 cells in transformed control wells (n=5), untransformed control wells (n=15), and *LIN28B* KD wells (n=18). (e) Proportion of B lymphoid (CD19+), myeloid (CD11b/CD14/CD1+) and NK (CD56+) cells assessed by flow cytometry from transformed control, untransformed control and *LIN28B* KD wells at day 21. (Data shown as % of viable hCD45 cells, mean+SEM).

### *LIN28B+* ^CRISPR*-KMT2A::AFF1*^ALL has a more proliferative and aggressive phenotype

Analysis of publicly available datasets showed that *LIN28B* expression is heterogeneous both in *KMT2A::AFF1* iALL patients (36, 58) and in the ^CRISPR*-KMT2A::AFF1*^ALL model (Figure 4a). Comparison of the gene expression profiles of high (*LIN28B+*, upper quartile of normalized counts) and low (LIN28B-, lower quartile: normalised counts=0) *LIN28B* expressing leukemias in primary patients and ^CRISPR*-KMT2A::AFF1*^ALL model, showed a significantly lower expression of *IGF2BP1* in the *LIN28B-* group (p=<0.0001) (Figure 4b). Gene set enrichment analysis demonstrated downregulation of MYC targets and mTORC1 signalling in the *LIN28B-* CRISPR-KMT2A::AFF1ALL (Figure 4c). To explore the significance of *LIN28B* expression, we compared survival curves of the primary *LIN28B+ and LIN28B-* ^CRISPR*-KMT2A::AFF1*^ALL mice. Leukemia latency in *LIN28B+* ^CRISPR-*KMT2A::AFF1*^ALL mice was significantly reduced (median survival 18.4 weeks) compared to *LIN28B-* ^CRISPR*-KMT2A::AFF1*^ALL mice (26.9 weeks; p=0.024) (Fig 4d). Similarly, *LIN28B+* ^CRISPR*-KMT2A::AFF1*^ALL blasts (n=3) showed significantly greater proliferation on MS5 stroma than LIN28B-^CRISPR*-KMT2A::AFF1*^ALL blasts (n=4) (p=0.026) (Figure 4e). To confirm that high *LIN28B* expression was driving this aggressive phenotype, *LIN28B* was knocked down in *LIN28B+* ^CRISPR*-KMT2A::AFF1*^ALL (n=3, Supplementary figure 3a). *LIN28B KD* ^CRISPR*-KMT2A::AFF1*^ALL blasts had significantly reduced proliferation compared with their matched controls in vitro (p=0.018) (Figure 4f). In vivo xenotransplantation assays using 35,000-50,000 cells/mouse, showed that peripheral blood engraftment kinetics was consistently faster in control ^CRISPR*-KMT2A::AFF1*^ALL transplanted mice compared to its paired *LIN28B* KD ^CRISPR-*KMT2A::AFF1*^ALL cohort (Suppl Fig 3b). Despite sample-to-sample variability in leukemia engraftment (Suppl Fig 3c), overall there was a significantly shorter median latency of leukemia development in mice transplanted with control ^CRISPR*-KMT2A::AFF1*^ALL (64 days) versus mice transplanted with *LIN28B* KD ^CRISPR*-KMT2A::AFF1*^ALL (78 days) (p=0.035) (Figure 4g). These data demonstrate that although LIN28B is expressed at high levels in only a subset of iALL patients and ^CRISPR*-KMT2A::AFF1*^ALL, it is important for maintaining a proliferative phenotype *in vivo* and *in vitro*.

**Figure 4.**
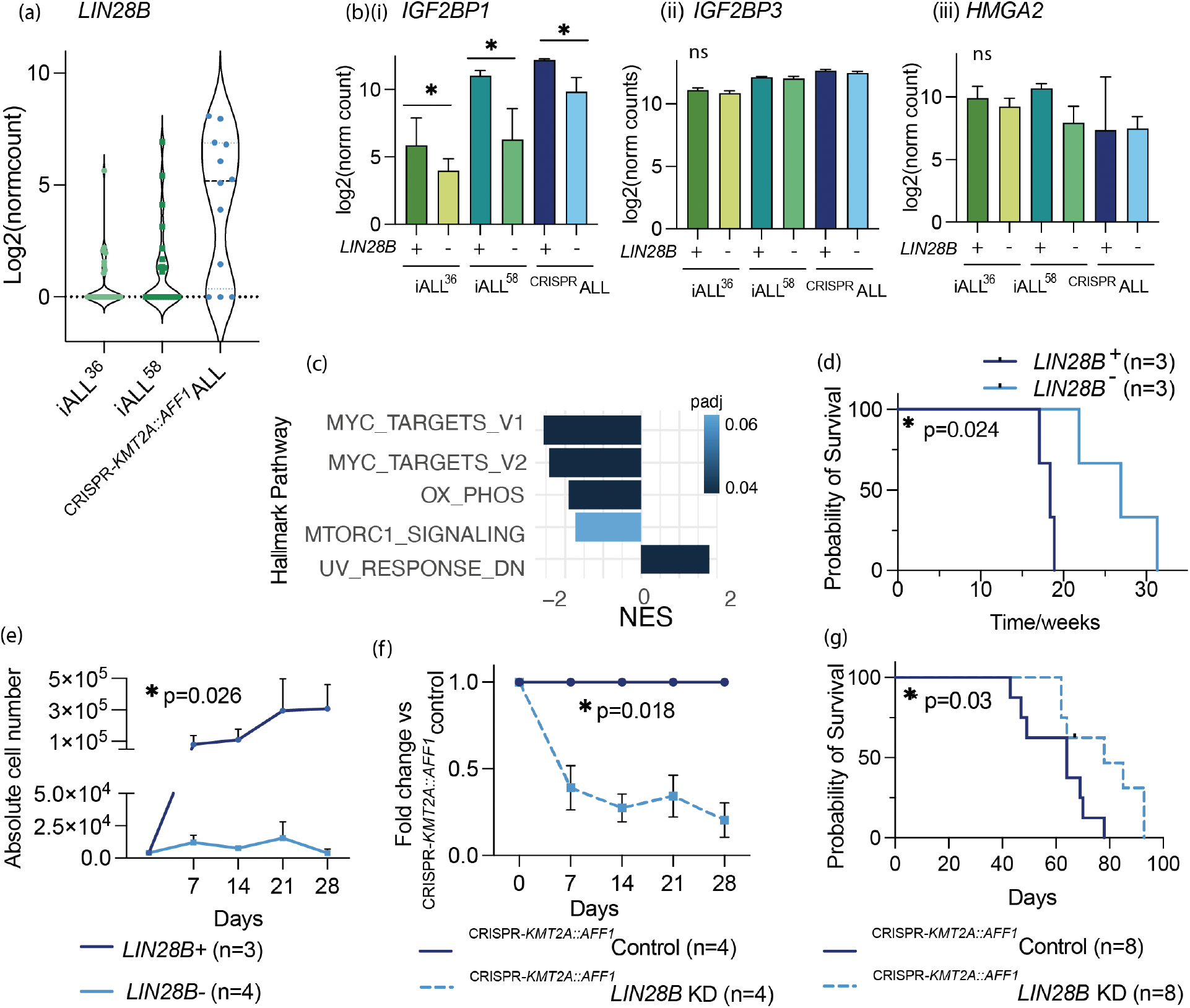
(a) log2 normalised expression of *LIN28B* in two *KMT2A:AFF1* iALL datasets (n=35)(36), (n=19)(58); and ^CRISPR-*KMT2A::AFF1*^ALL (n=12). (b) Comparison of *LIN28B+* and *LIN28B-KMT2A:AFF1* leukemias: iALL dataset 1(36) LIN28B+ (n=9), LIN28B-(n=26); iALL dataset 2(58) LIN28B+ (n=5), LIN28B-(n=9); ^CRISPR^ALL LIN28B+ (n=3), ^CRISPR^ALL LIN28B-(n=3), for expression of LIN28B pathway genes: (i) *IGF2BP1*, (ii) *IGF2BP3* (iii) *HMGA2* (significance based on one way ANOVA). (c) GSEA for Hallmark pathways up and downregulated in LIN28B-^CRISPR*-KMT2A::AFF1*^ALL compared to LIN28B+ ^CRISPR*-KMT2A::AFF1*^ALL. (d) Survival curves for primary ^CRISPR*-KMT2A::AFF1*^ALL xenografts LIN28B^+^ (n=3) or LIN28B^-^ (n=3). (e) Proliferation of *LIN28B+* (n=3) and *LIN28B-* ^CRISPR*-KMT2A::AFF1*^ALL blasts (n=4) on MS5 cocultures over 28 days, absolute cell number depicts viable, hCD45+ cells at each timepoint per well. (f) Proliferation of *LIN28B* KD ^CRISPR*-KMT2A::AFF1*^ALL blasts on MS5 stroma viewed as fold change in cell number (viable, hCD45+ cells) relative to matched ^CRISPR-KMT2A-AFF1^ALL control leukemia. (g) Survival curve for ^CRISPR*-KMT2A::AFF1*^leukemia (n=3 biological samples, transduced with scr control (n=8 mice) or *LIN28B KD* (n=8 mice)).

### mRNA binding partners of LIN28B

To investigate the mechanism through which LIN28B (an RNA binding protein) alters the molecular and biological characteristics of *KMT2A::AFF1* iALL, we used SEM cells, a well-established *KMT2A::AFF1* B-ALL cell line model in order to obtain sufficient RNA. LIN28B acts both via let 7 miRNA-dependent and let-7 independent mechanisms. To interrogate let-7 independent functions of LIN28B, RNA Immunoprecipitation sequencing (RIPseq) was performed to identify RNAs bound to LIN28B and therefore more likely to be directly regulated by it in SEM cells. Similarly to ^CRISPR*-KMT2A::AFF1*^ALL, *LIN28B* KD in SEM cells significantly impaired proliferation in liquid culture (p=0.01) via apoptosis (Suppl Figure 4a-c) and colony formation in methylcellulose compared to control cells (Supp Fig 4d). *LIN28B* KD SEM cells also demonstrated increased leukemia latency *in vivo* (p=<0.0001) (Suppl Figure 4e). As in FL CD34+ cells, *LIN28B* KD led to downregulation of G2M gene sets, E2F targets and MYC targets (Figure 5a). As anticipated, let 7 mirRNA target gene sets were also downregulated upon *LIN28B* KD (Suppl Fig 5a). Together, these data show that *LIN28B* is essential for SEM cell survival and that it regulates many of the same pathways as in primary cells making it an appropriate model to interrogate mechanistic pathways.

**Figure 5.**
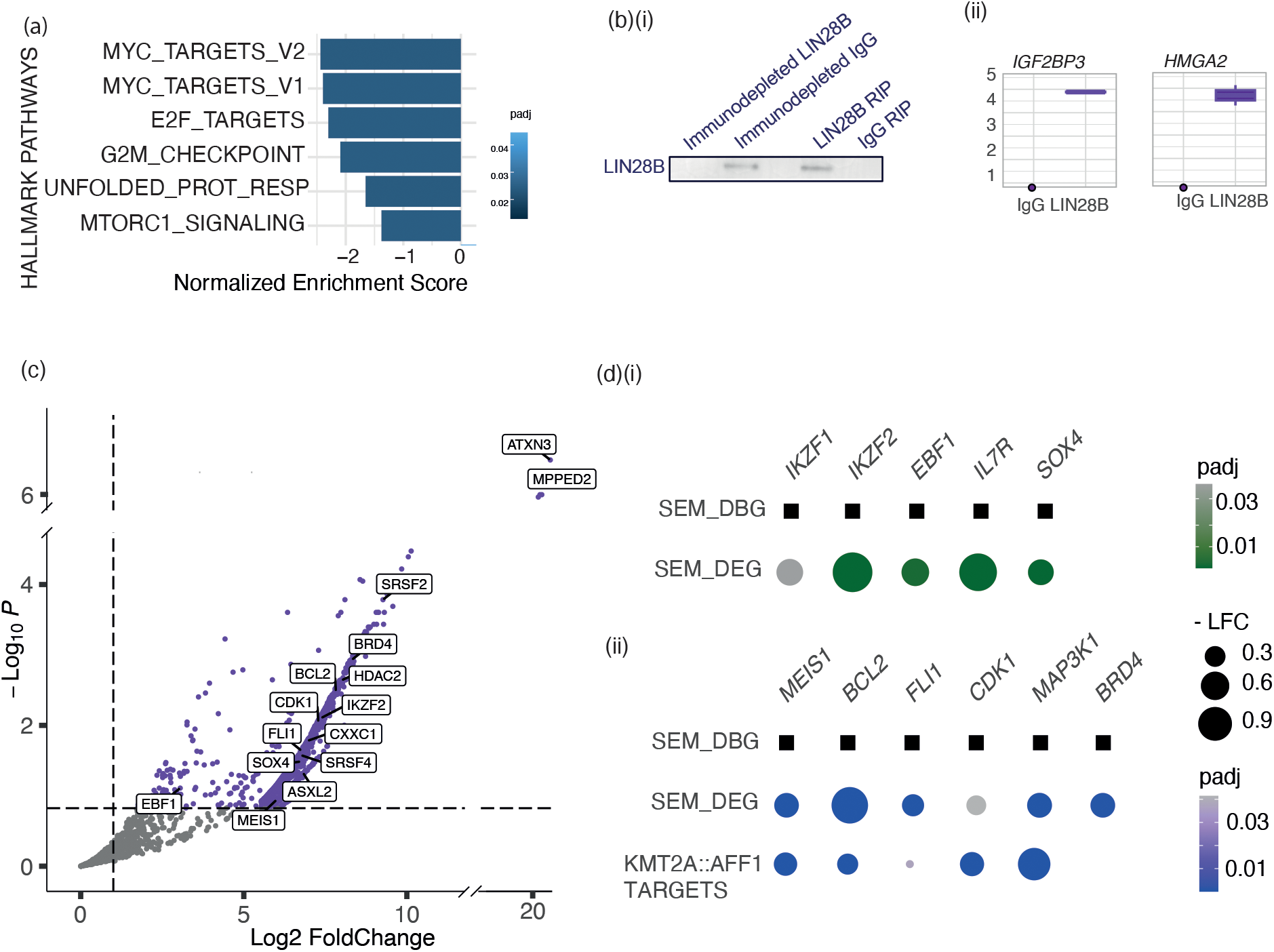
(a) Gene set enrichment analysis showing pathways downregulated in LIN28B KD SEM cells compared to control SEM cells (b)(i) Representative western blot of the immunodepleted fraction and immunoprecipitated (RIP) fraction of LIN28B or control IgG immunoprecipitation (ii) Log2 normalised counts for *IGF3BP3* and *HMGA2*, dissociated from immunoprecipitated LIN28B in RIPseq with LIN28B (n=3) or IgG (n=2). (c) Volcano plot of genes bound by LIN28B identified from DESeq2 analysis of LIN28B and IgG bound genes (significant genes coloured in purple: log2FC>0.5, padj<0.1). Labelled are those transcripts, that are also downregulated upon *LIN28B* KD in SEM and FL CD34+ cells. (d) Examples of (i) B-lymphoid genes and (ii) KMT2A::AFF1 target genes identified by KMT2A::AFF1 knockdown in SEM cells(59), that are both bound by LIN28B and downregulated upon *LIN28B* KD in SEM cells (log2FC<-0.2, padj<0.1, in all datasets).

RIPseq was performed using antibodies specific to LIN28B and results were compared to an IgG control immunoprecipitation. Successful immunoprecipitation of LIN28B was confirmed by western blot (Figure 5b(i)). Raw feature counts for genes were filtered, to exclude genes with zero counts, or zero counts in two of the three LIN28B replicates, prior to DEseq2 based analysis. Identification of known targets such as *IGF2BP3* and *HMGA2* that were enriched in the LIN28B RIP compared to the IgG control validated the selectivity of the experiment (Figure 5b(ii)). Overall, there were 1620 differentially bound genes (log2FC>0.5, padj <0.1) enriched in the LIN28B RIP versus the IgG control. To identify LIN28B bound mRNAs that were stabilised by LIN28B, datasets of bound RNA were intersected with DEGs from RNAseq datasets for SEM *LIN28B* KD vs SEM control cells. This identified 370 genes that were both bound by LIN28B and downregulated in SEM *LIN28B* KD cells and pathway analysis of these genes again enriched for G2M, E2F, MYC, targets and cell cycle genes (Suppl Fig 5b).

### B lymphoid genes bound by LIN28B

As in FL CD34+ cells (Fig 2c), *LIN28B* KD in SEM cells caused downregulation of B-lymphoid genes (NES = -1.8, padj 0.0096) (Suppl Fig. 5c). Key lymphoid genes bound by LIN28B and downregulated upon *LIN28B* KD in SEM cells include, *EBF1, LEF1, IL7R*, and *SOX4* (Figure 5d(i)). *IKZF1* and *IKZF2* were also downregulated in *LIN28B* KD FL CD34+ cells. This suggests that LIN28B binds to and stabilises B-lymphoid genes in both normal and malignant cells.

### KMT2A::AFF1 target genes bound by LIN28B

To investigate whether LIN28B is essential for the regulation of key KMT2A::AFF1 target genes that drive leukemogenesis, we intersected known KMT2A::AFF1 target genes established from published nascent RNAseq following KMT2A::AFF1 KD *(59)* with our RIPseq data. Of the 370 LIN28B bound genes which are downregulated upon *LIN28B* KD, almost a quarter (88) are KMT2A::AFF1 targets *(59)*. These include important downstream targets *MEIS1, BCL2*, and *FLI1*, which are also downregulated upon *LIN28B KD* in SEM and FL CD34+ cells. Further genes which enhance proliferation and are KMT2A::AFF1 targets, for example CDK1 and MAP3K1 were also stabilised by LIN28B (Figure 5 d(ii)).

In addition to KMT2A::AFF1 target genes, several chromatin proteins known to be essential for *KMT2Ar* leukemogenesis were also regulated by LIN28B. For example, BRD4 is highly bound by LIN28B and downregulated upon *LIN28B* KD in SEM cells and FL CD34+. BRD4 has been shown to promote transcription in KMT2A::AFF1 cells(60) and to be essential for survival of *KMT2Ar* leukemias(61, 62).

Overall, we find that LIN28B binds and regulates a common set of gene targets across different cell types but it also has cell type specific activity. LIN28B specifically regulates genes essential for normal fetal B lymphopoiesis, which may also contribute to *KMT2A::AFF1* B-ALL initiation.

## DISCUSSION

iALL is a poor prognosis leukemia that originates *in utero* and displays a fetal specific molecular profile(40). *LIN28B*, a fetal specific gene encoding an RNA binding protein (RBP), is aberrantly expressed in many cancers(45, 46, 48), including those with putative fetal origins(47) such as iALL. While it is known that human fetal B-lymphopoiesis is distinct from postnatal B-lymphopoiesis(35, 63, 64), and that *Lin28b* is important for murine fetal B-lymphopoiesis(52, 53), the exact role of *LIN28B* in human fetal B-cell development and/or promoting initiation and maintenance of iALL is not clear. A human FL-derived ^CRISPR*-KMT2A::AFF1*^ALL model, allows us to interrogate the role of LIN28B in the initiation and maintenance of iALL in the right cellular context.

*LIN28B* and its partner genes have highest expression in primitive human fetal HSPC, with a downregulation as B-cell commitment occurs. In human FL CD34+ cells, *LIN28B* is required for B- and NK-lymphopoiesis and also initiation of *KMT2A::AFF1* ALL. In leukemias where it is expressed, *LIN28B* confers a more aggressive phenotype. Together, our results show that *LIN28B* has a dual role in maintaining normal fetal lymphopoiesis as well as providing a precondition for the development of aggressive infant leukemia.

The mechanism of action of LIN28B is likely to be multifactorial. It specifically binds to and regulates genes essential for normal B-lymphopoiesis, which may also contribute to *KMT2A::AFF1* B-ALL initiation. In addition, LIN28B is needed to maintain mRNA stability of important oncogenes, including those regulated by KMT2A::AFF1 such as *MEIS1* and *BCL2* which probably also contributes to its essentiality for leukemia initiation and maintenance. The synergy of driving a B-lymphoid programme and stabilisation of oncogenes by LIN28B, provides the permissive context in human fetal cells for initiation of B-ALL rather than AML(65).

These data provide the first evidence of a role for *LIN28B* in human fetal B-lymphopoiesis and infant leukemia initiation, and highlights possible mechanisms that could be targeted in this aggressive disease.

## ACKNOWLEDGEMENTS

We are grateful for the technical support received from the WIMM Flow Cytometry Core, WIMM Sequencing Core, WIMM Centre for Computational Biology, MRC WIMM Virus screening facility and John Radcliffe Biomedical Sciences Department. This work was funded by a DPhil in Cancer Sciences studentship awarded to RL by Cancer Research UK (Oxford Cancer Centre). TAM and AS are both supported by the Medical Research Council (MRC, UK) Molecular Haematology Unit grant MC_UU_00029/6. AR is supported by a Wellcome Trust Clinical Research Career Development Fellowship (216632/Z/19/Z) and Medical Research Council (MRC, UK) Molecular Haematology Unit grant MC_UU_00029/7. The human fetal material was provided by the Joint MRC/Wellcome Trust Grant 099175/Z/ 12/Z Human Developmental Biology Resource (http://hdbr.org).

## Notes

### Competing Interest Statement

T.A.M. is a paid consultant for and shareholder in Dark Blue Therapeutics Ltd.
B.P. is a co-founder and shareholder in Alethiomics and provides consultancy for Incyte, BluePrint Medicine and Novartis.

