## Supplemental Material for "The fetal specific gene *LIN28B* is essential for human fetal B-lymphopoiesis and initiation of KMT2A::AFF1 infant leukemia"

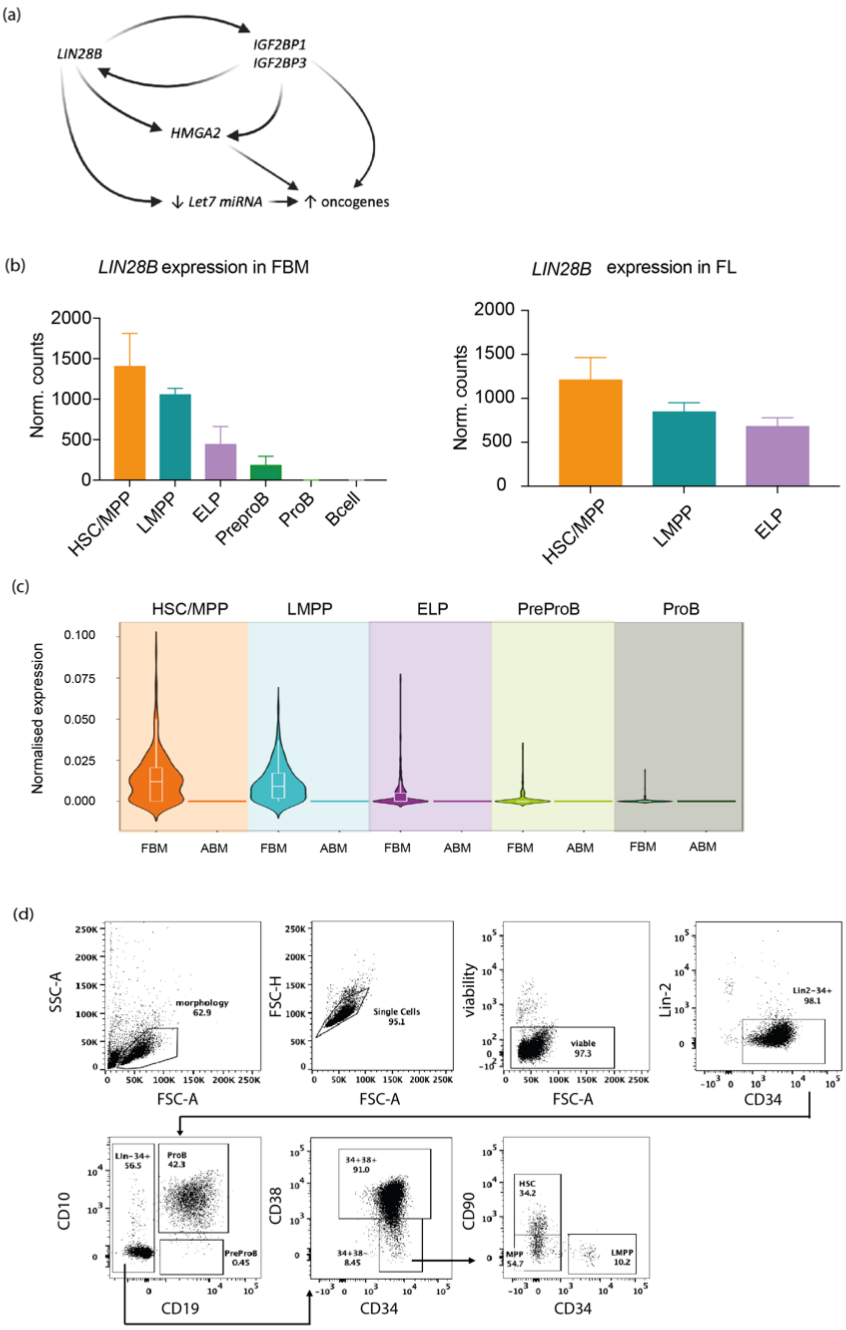


**Suppl Fig 1** (a) Schematic of the LIN28B pathway. LIN28B stabilises IGF2BP1 /IGF2BP3, both enhance *HMGA2* expression. LIN28B negatively regulates the tumor suppressors, let-7 miRNA. These result in increased expression of oncogenes (b) Bulk RNAseq data showing *LIN28B* gene expression in sorted fetal HSPC from FBM (left, n=3) and FL (right, n=3). Data presented as mean normalised counts (HSC: hematopoietic stem cell; MPP: multipotent progenitor; LMPP: lymphoid primed MPP; ELP: early lymphoid progenitor) (c) Single cell qPCR data showing normalised expression of *LIN28B* in FBM and ABM HSPC (d) Representative flow plots from an ABM sample showing sorting strategy used to isolate HSC, MPP and LMPP for mini-bulk RNA-seq from FL, FBM and ABM CD34+ cells.

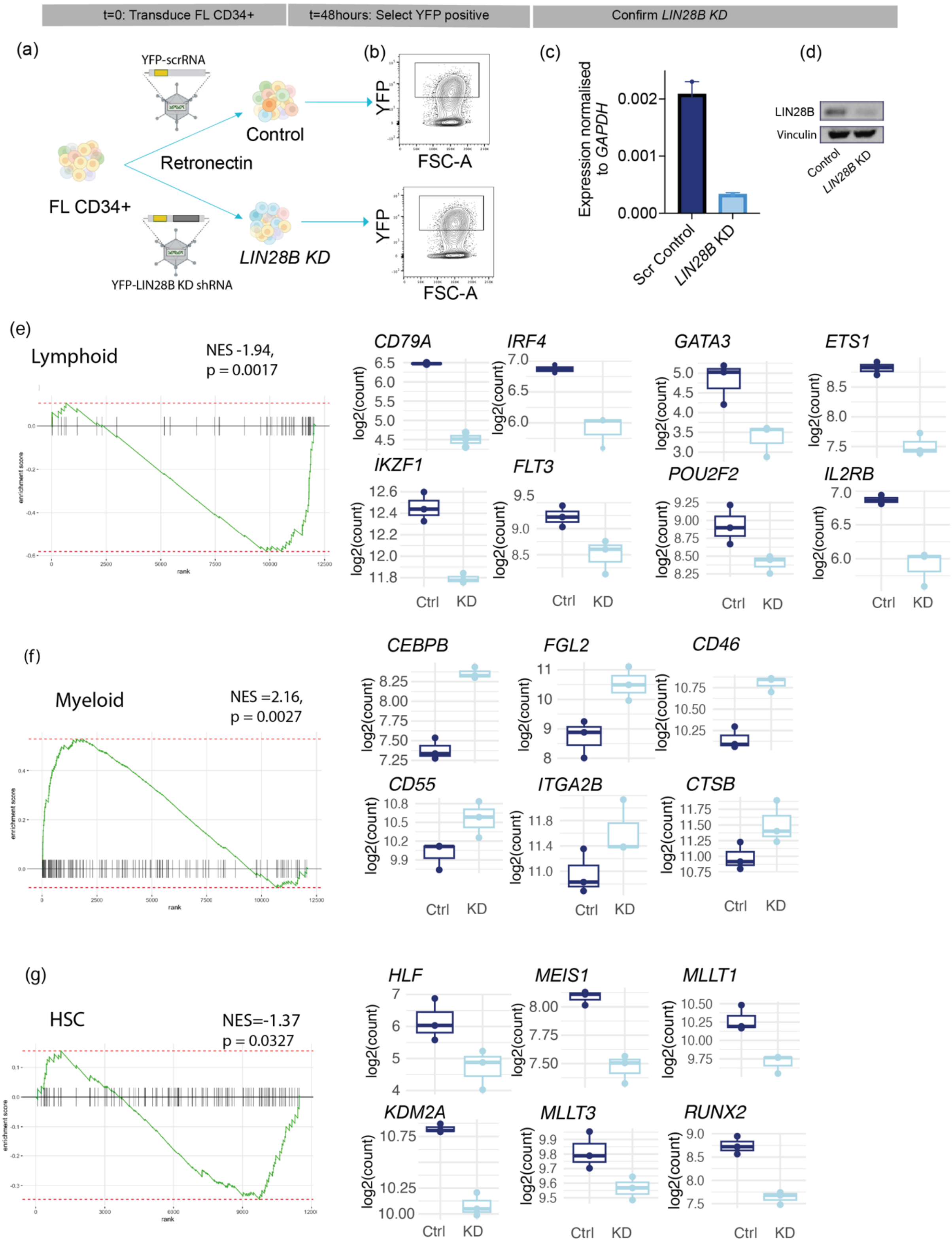


**Suppl Figure 2** (a) Schema for shRNA *LIN28B* KD in human FL CD34+ cells, followed by (b) FACS selection of YFP+ cells at 48 hours after transduction n=5. (c) qPCR for *LIN28B* expression of control and *LIN28B* KD FL CD34+ cells. (d) Representative Western blot for LIN28B and vinculin expression in control and LIN28B KD FL CD34+ cells at 48 hours after transduction. Gene set enrichment analysis of pathways affected by *LIN28B* KD in FL CD34+ cells (e) fetal lymphoid geneset(1) (f) fetal myeloid geneset(1) and (g) cord blood HSC geneset(2). Box plots show log2(normalised counts) for control and *LIN28B* KD FL CD34 cells for selected lymphoid, myeloid and HSC genes.

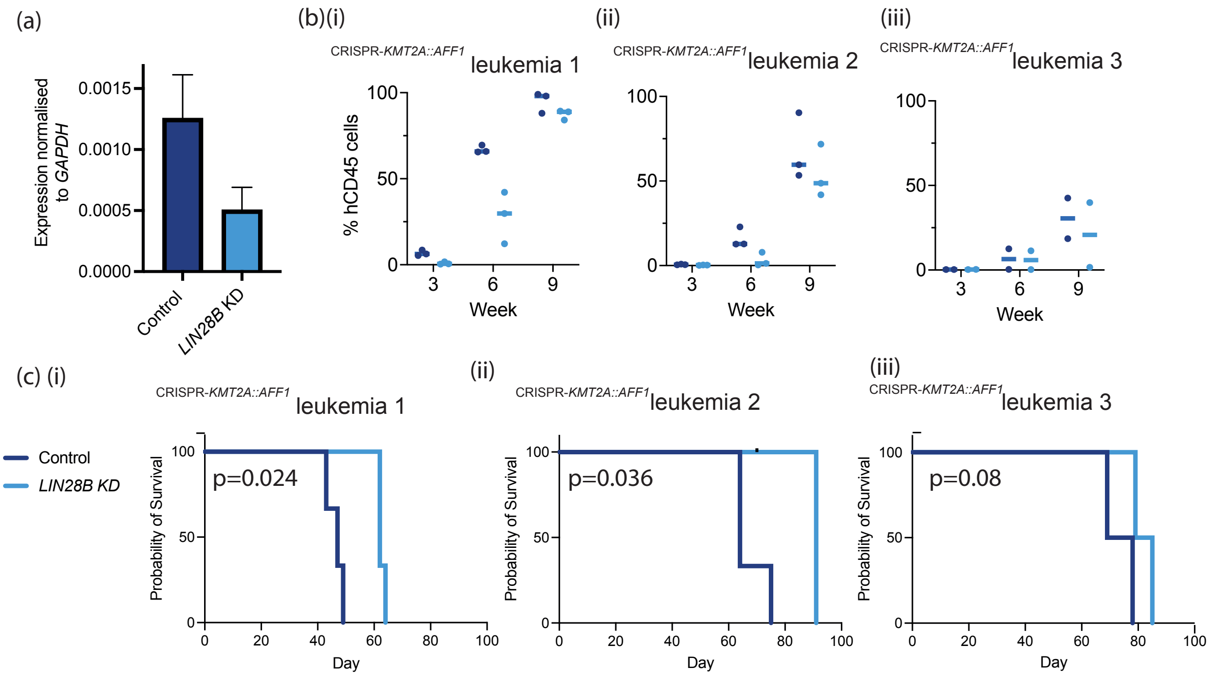


**Suppl Figure 3** (a) qPCR for *LIN28B* expression of control and *LIN28B* KD $^{\text{CRISPR-}\textit{KMT2A::AFF1}}$ leukaemia: analysis done on YFP+ cells isolated by FACS at 48hours after transduction. (b) Proportion of viable human CD45 (hCD45) cells detectable in the blood of mice transplanted with 35,000-50,000 $^{\text{CRISPR-}\textit{KMT2A::AFF1}}$control (dark blue) or $^{\text{CRISPR-}\textit{KMT2A::AFF1}}$*LIN28B* KD (light blue) cells, measured at 3 weekly intervals from three different $^{\text{CRISPR-}\textit{KMT2A::AFF1}}$ALL samples (i-iii). (c) Survival curves for mice injected with $^{\text{CRISPR-}\textit{KMT2A::AFF1}}$control or $^{\text{CRISPR-}\textit{KMT2A::AFF1}}$*LIN28B* KD cells as described in (b).

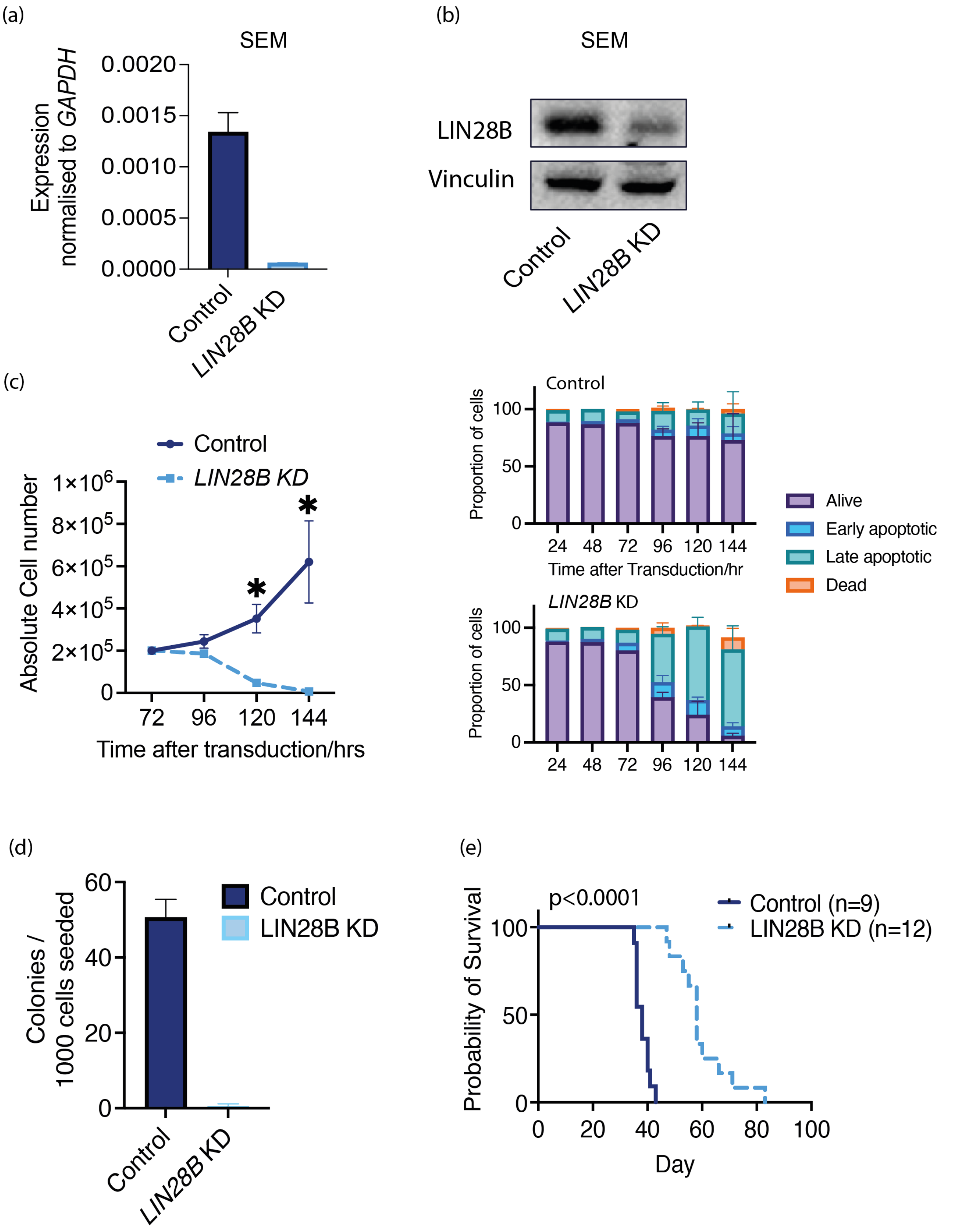


**Suppl Figure 4** (a) qPCR for LIN28B expression of control and *LIN28B* KD SEM cells isolated by FACS 72 hours after transduction n=5. (b) Representative Western blot for LIN28B and vinculin expression in control SEM and *LIN28B* KD SEM cells at 72 hours after transduction (c) Left: Proliferation of control SEM and *LIN28B* KD SEM cells in liquid culture monitored at 12 hourly intervals; Right: proportion of live, early apoptotic, late apoptotic and dead cells measured over 6 days in control SEM and *LIN28B* KD SEM cells in liquid culture, n=3. (d) Methylcellulose colony forming assay showing no. of colonies generated per 1000 cells plated for control and *LIN28B KD* cells at day 14, p<0.001, n=5. (e) Survival curves for mice injected with 1M control SEM or *LIN28B* KD SEM cells.

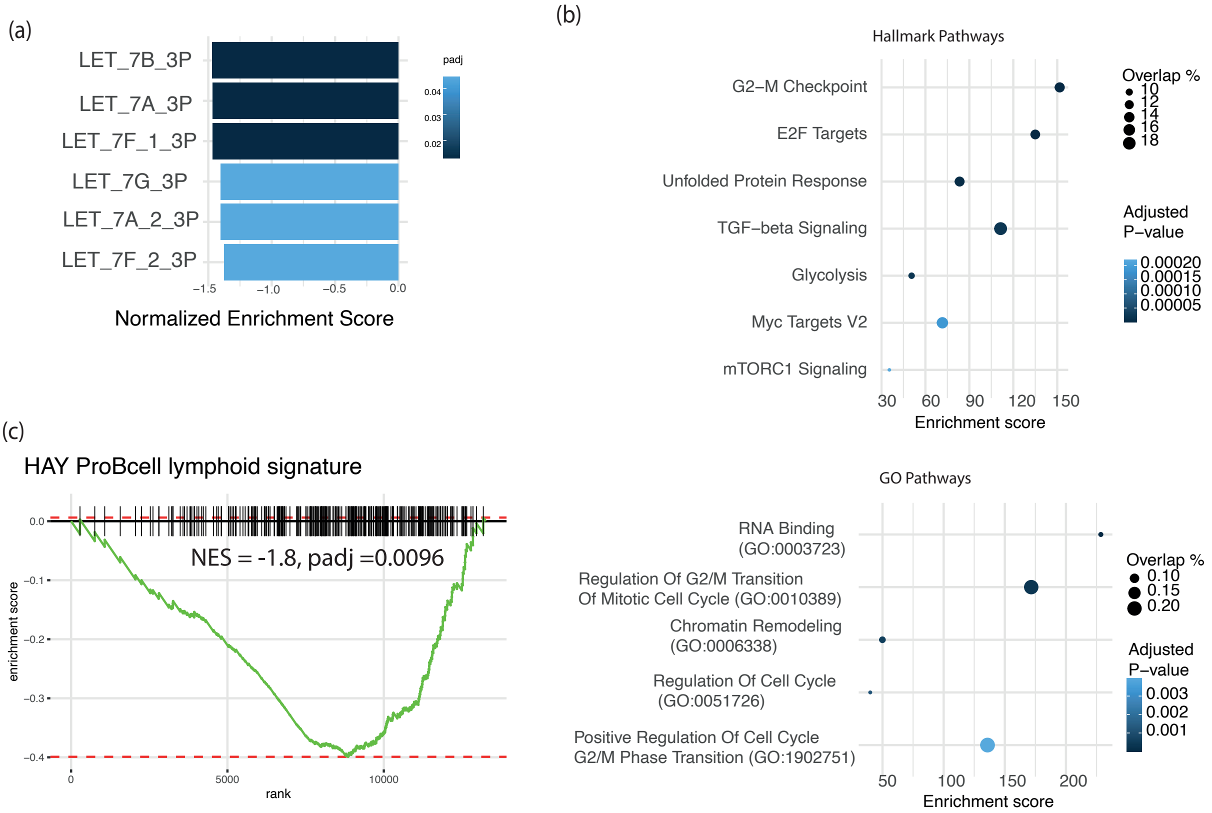


**Suppl Figure 5** (a) GSEA of let 7 miRNA target genesets comparing *LIN28B* KD SEM cells (n=3) to control SEM cells (n=3). (b) Bubble plot of enriched genesets for LIN28B bound genes in SEM cells (c). GSEA showing downregulation of a ProB lymphoid gene signature(3) in LIN28B KD SEM cells (n=3) compared to control SEM cells (n=3).

**Supplementary Table 1: LIN28B pathway: Differentially expressed genes in HSC, MPP and LMPP between fetal and adult**

| gene | Tissue compared to ABM | HSPC | log2FC | padj |
|---|---|---|---|---|
| LIN28B | FL | HSC | 12.70 | 2.36E−21 |
| | FBM | HSC | 11.56 | 0.000708 |
| | FL | MPP | 11.09 | 8.9316E-09 |
| | FBM | MPP | 12.27 | 1.51E−12 |
| | FL | LMPP | 11.82 | 4.06E−61 |
| | FBM | LMPP | 11.76 | 1.17E−27 |
| IGF2BP1 | FL | HSC | 13.00 | 2.64E−02 |
| | FBM | HSC | 10.75 | 0.000416 |
| | FL | MPP | 11.11 | 8.0181E-12 |
| | FBM | MPP | 11.35 | 3.55E−06 |
| | FL | LMPP | 12.01 | 5.3E−128 |
| | FBM | LMPP | 11.01 | 1.57E−26 |
| IGF2BP3 | FL | HSC | | ns |
| | FBM | HSC | | ns |
| | FL | MPP | | ns |
| | FBM | MPP | | ns |
| | FL | LMPP | 3.23 | 0.023935 |
| | FBM | LMPP | | ns |
| HMGA2 | FL | HSC | 1.81 | 6.86E−06 |
| | FBM | HSC | 3.19 | 0.002833 |
| | FL | MPP | 1.50 | 0.0501 |
| | FBM | MPP | 2.20 | 0.027659 |
| | FL | LMPP | | ns |
| | FBM | LMPP | | ns |

**Supplementary Table 2: Taqman Assays**

| Gene | Company | Catalogue Number |
|---|---|---|
| *LIN28B* | Thermofisher | Hs01013729_m1 |
| *GAPDH* | Thermofisher | Hs02758991_m1 |

**Supplementary Table 3: Primers for ddPCR**

| Primer | Sequence |
|---|---|
| KMT2A fwd | GAAAATCGCTTGAACTTTGG |
| AFF1 rev | CTCCCCTGCTCAAAATTATC |
| RPP30 fwd | GATTTGGACCTGCGAGCG |
| RPP30 rev | GCGGCTGTCTCCACAAGT |

**Supplementary Table 4: Probes for ddPCR**

| Probe | Sequence |
|---|---|
| AFF1 probe | [6FAM]ACCATGGAGGCTTTGTTGGC |
| RPP30 probe | [HEX]CTGACCTGAAGGCTCT |

**Supplementary Table 5: Antibodies for Western Blot and RIPseq**

| Protein | Company | Catalogue number | Application |
|---|---|---|---|
| LIN28B | Cell signalling | 4196 | Western blot/RIPseq |
| Vinculin | Cell signalling | 4650 | Western blot |
| IgG | Cell signalling | 3900 | RIPseq |

**Supplementary Table 6: Antibodies for flow cytometry/FACS**

| Antibody | Fluorphore | Clone | Company | Catalogue Number |
|---|---|---|---|---|
| CD3 | BV711 | OKT3 | Biolegend | 317328 |
| CD3 | PerCPcy5.5 | OKT3 | Biolegend | 317335 |
| CD10 | PE-Cy7 | ebioCCB-CALLA | Life Technologies | 11-0106-42 |
| CD11b | PerCP-Cy5.5 | ICRF44 | BioLegend | 301328 |
| CD14 | PerCP-Cy5.5 | M5E2 | Biolegend | 301824 |
| CD16 | PerCP-Cy5.5 | 3G8 | BioLegend | 302028 |
| CD19 | APC | HIB19 | BioLegend | 302212 |
| CD33 | FITC | P67.6 | BioLegend | 366620 |
| CD34 | BV421 | 561 | BioLegend | 343610 |
| CD45 | AF700 | 2D-1 | Life Technologies | 56-9459-42 |
| CD56 | BV605 | HCD56 | Biolegend | 318334 |
| CD56 | PerCP-Cy5.5 | HCD56 | Biolegend | 318322 |
| CD90 | BV421 | 5.00E+10 | BioLegend | 328122 |
| CD133 | PE | AC133 | Miltenyi | 130-113-108 |
| CD235a | PerCP cy5.5 | HIR2 | BioLegend | 306614 |
| mouse CD45.1 | APC-Cy7 | 30-F11 | BioLegend | 103116 |
| Viability | ef506 | | Life Technologies | 65-0866-18 |
| Brilliant stain buffer plus | | | BD Biosciences | 566385 |

### METHODS

#### Cell culture

**FL CD34+ and $^{\text{CRISPR-}\mathit{KMT2A::AFF1}}$ leukemia co-culture:** Confluent MS5 stromal layers were prepared as described previously. 4000 cells were seeded to MS5 layers. FL CD34+ cells were maintained in alphaMEM in 10% heat inactivated fetal bovine serum (HI-FBS), 100 U/ml penicillin, 100 µg/ml streptomycin, 2 mM L-glutamine, 50 µM 2-mercaptoethanol, 10 mM HEPES, SCF 20ng/ml, FLT3L 10ng/ml, IL-2 10ng/ml and IL-7 5ng/ml (Peprotech). $^{\text{CRISPR-}\mathit{KMT2A::AFF1}}$ leukemia were maintained on MS5 stromal cells in SFEM II supplemented with 10% HI-FBS, 2 mM L-glutamine, IL-3 10ng/ml and IL-7 5ng/ml (Peprotech). All cocultures were incubated at 37°C in 5% CO2 with twice-weekly half-volume medium changes. Flow-cytometric readouts were performed weekly. Cells were replated on fresh MS5 stromal cells every 2 weeks.

**SEM cell culture:** cells were cultured in Iscove's modified Dulbecco's medium (IMDM) with 10% HI - FBS and 2 mM L-glutamine, the cell density maintained between 0.5-4 *$10^{6}$cells/mL.

#### Lentiviral transduction:

*LIN28B shRNA* (sense strand: CCGGGCACCAGAAGAGCAAAGCAAACTCGAGTTTGCTTTGCTCTTC TGGTGCTTTTTG) or a scrambled (scr) RNA control was cloned into pLKO.eYFP plasmid. High titre lentivirus (LV) was generated from pLKO.eYFP plasmid (EV control), pLKO.eYFP + scr RNA (scr control) or pLKO.eYFP + LIN28B shRNA (*LIN28B* KD) by MRC WIMM Virus screening facility.

Due to limitations in the availability of primary xenograft material $^{\text{CRISPR-}\mathit{KMT2A::AFF1}}$ leukemia co-cultures were performed on blasts derived from the BM of secondary xenografts using the established thresholds defined in primary LIN28B+ or LIN28B- $^{\text{CRISPR-}\mathit{KMT2A::AFF1}}$ leukemia. FL CD34+ and $^{\text{CRISPR-}\mathit{KMT2A::AFF1}}$ leukemia were transduced either scr control LV or *LIN28B* KD LV using retronectin enhanced protocol from the manufacturer, Takara Bio. Transduced cells were isolated at 48 hours based on viable cells with high YFP expression by FACS. SEM cells were transduced by spinoculation with EV control LV or *LIN28B* KD LV, transduced cells were isolated by FACS based on viable cells with high YFP at 72 hours.

**CRISPR-Cas9 editing of LV transduced cells:** LV transduction of FL CD34+ cells performed as described above. 24 hours after transduction, CRISPR-Cas9 editing was performed as described in (4), then edited cells returned to StemLine II media supplemented with SCF 100ng/ml, FLT3L 100ng/ml and TPO 100ng/ml. Viable cells with high YFP expression were isolated by FACS 48 hours after transduction.

**Xenotransplantation assays**

**Animals :** Experiments were performed under the Animal (Scientific Procedures) Act 1986 with institutional approval by local ethical review body. Experimental animals were 8–12-week-old NOD.Cg-Prkdc$^{scid}$ Il2rg$^{tm1Wjl}$/SzJ (NSG, Jackson Laboratories) mice.

**$^{\text{CRISPR-}\textit{KMT2A::AFF1}}$ leukemia generation**

$^{\text{CRISPR-}\textit{KMT2A::AFF1}}$ leukemia was generated from second trimester human FL HSPC as described in (4). Briefly, 8-12 week old female NSG mice were sub-lethally irradiated with 2 doses of 1.25Gy prior to tail vein injection with 3000-30 000, edited human FL HSPC (n=12).

***LIN28B* KD $^{\text{CRISPR-}\textit{KMT2A::AFF1}}$ALL xenotransplantation:** 35,000-50,000 leukemic blasts from BM of mice with $^{\text{CRISPR-}\textit{KMT2A::AFF1}}$ ALL were transplated via tail vein injection in sublethally irradiated NSG mice. (LIN28B KD (n=8) or scr control (n=8)) .

**SEM xenotransplantation:** 8–12-week-old male NSG mice received 1 million viable, SEM LIN28B KD (n=12) or control cells (n=9) by tail vein injection. Engraftment was monitored with peripheral blood sampling every 3 weeks. Mice were monitored for signs of ill-health and culled in the presence of weight loss >15%, limb paralysis, PB >80% hCD45 cells, or reduced movement, hunched posture and starry coat.

**Molecular assays**

**RT qPCR:** mRNA was extracted from cells using an RNeasy Micro Kit (Qiagen) and complementary DNA (cDNA) generated using the SuperScript II Kit (Invitrogen). cDNA was analyzed by Taqman qPCR, using the housekeeping gene *GAPDH* for gene expression normalization. Taqman probes were used for qPCR QuantStudio3 Real-Time PCR System (Thermo Fisher). A list of Taqman assays are provided in Supplementary Table 2.

**ddPCR:** Cells were lysed in SingleShot Cell Lysis Buffer with proteinase K. A ddPCR droplet generation mix generated from cell lysate, supermix for probes (Bio-Rad), bespoke probes and primers designed for KMT2A, AFF1 and reference gene RPP30 (Supplementary Table 3 and 4), CviQI restriction enzyme and PCR grade water. Droplets are generated using QX200 AutoDG loaded with Oil for Probes. PCR reaction on themocycle (Bio-Rad) with cycling conditions: 95 °C for 10 min, 40 cycles at 94 °C for 30 s, 53 °C for 75 s, 98 °C for 10 min, and 12 ℃ for 5 min. PCR plate containing PCR amplified droplets read on QX200 Droplet Reader run for Rare Event Detection and analyzed using the Quanta Soft software (Bio-Rad).

**Western blot:** Cells were lysed in 2X NuPAGE LDS Sample Buffer (INvitrogen). Samples were heated at 95°C for 10 minutes and sonicated using a Diagenode Bioruptor. Western blotting were previously described in (5).Antibodies used d are detailed in Supplementary Table 5.

**Methylcellulose-based colony-forming unit assays:** Methocult media was made by addition of IMDM, FBS, 200mM L-glutamine, and 100 U/ml penicillin, 100 µg/ml streptomycin,to methocult H4100 (Stemcell Technologies). SEM cells (EV control (n=5) or LIN28B KD (n=5)), were added to methocult, seeding density 500-1000 cells/plate. Cells in methocult media were cultured at 37deg c in 5% for 14 days and colony number assessed.

**Flow cytometry:** Cells were stained with fluorophore-conjugated monoclonal antibodies in phosphate-buffered saline with 2% FBS and 1 mM EDTA for 30 min and analyzed using BD Fortessa X20 or X50 instruments. Antibodies used are detailed in Supplementary Table 6. Analysis was performed using FlowJo 10 software.

**NGS**

**Bulk RNAseq:** 160 000-300 000 viable, hCD45+, CD19+ $^{\text{CRISPR-}\textit{KMT2A::AFF1}}$ leukemia were isolated from BM leukaemic mice. Starting cell numbers were 100 000 cells for viable, FL CD34+ (scr control and LIN28B KD) and 1 million viable, SEM (EV control and LIN28B KD) cells, both these were performed in triplicate for each experimental condition. Cells were lysed and total RNA was extracted using an RNeasy Microi Kit(Qiagen). Library preparation and sequencing were performed as previously described (4). RNA-seq protocols for low cell number (100 sorted cells) subpopulations of ABM HSPC was performed in triplicate as described in (6).

**RIPseq**: RIPseq was performed on 100 million SEM cells using Magna RIP RNA Binding Protein Immunoprecipitation Kit (Merck). Immunoprecipitation of LIN28B (n=3) or control IgG (n=2) performed using antibodies described in (Suppl Table 5). Western Blot confirmed LIN28B pull down and compared to immunodepleted fraction. The RNA extracted was purified using Monarch RNA cleanupkit (NEB) according to manufacturer's instructions. Library preparation and sequencing were performed as previously described(4).

**NGS analysis**:

**RNAseq:** Following RNAsequencing, the SeqNado pipeline (https://github.com/alsmith151/SeqNado), was used to assess read quality with the fastQC package, perform trimming using Trim Galore and map trimmed reads to the human genome assembly(hg38) using STAR. Protein coding reads were selected for downstream analysis. DEseq2 batch correction was applied and differentially expressed genes identified. For $^{\text{CRISPR-}\mathit{KMT2A::AFF1}}$ leukemia and iALL sample, LIN28B+ sample defined as those with expression in the upper quartile. LIN28B- had normalised counts =0.

**RIP-seq:** As for bulk RNA-seq, SeqNado pipeline was used to quality control, trim and map reads. Reads were filtered as for RNAseq, additionally only reads present in 2 of 3 of the LIN28B replicates were included. DEseq2 comparision of IgG and LIN28B RIP generated differentially expressed reads, p-adjusted filter $p<0.1$ applied. Gene set enrichment analysis performed using fsgea package (7), and pathway analysis: Enrichr portal (8).

**Statistical analysis:** To compare experimental groups, Student two sided t tests and analysis of variance followed by multiple comparison testing were used (as indicated in the figure legends). Log-rank(Mantel-Cox) test was applied to in vivo survival curves. Statistical analyses were performed by using GraphPad Prism v10.0.2 or R version 4.2.1 Data are expressed as mean ± standard error of the mean (SEM) unless otherwise indicated.
